# Oral Delivery of Kidney Targeting Nanotherapeutics for Polycystic Kidney Disease

**DOI:** 10.1101/2022.10.18.512444

**Authors:** Yi Huang, Jonathan Wang, Deborah Chin, Valeria Mancino, Jessica Pham, Hui Li, Kairui Jiang, Aparna Ram, Christopher Poon, Pei-Yin Ho, Georgina Gyarmati, János Peti-Peterdi, Kenneth R. Hallows, Eun Ji Chung

**Author notes:** Corresponding author: Eun Ji Chung University of Southern California Department of Biomedical Engineering 1002 Childs Way, MCB 357, Los Angeles 90089, CA, USA.

## Abstract

Autosomal dominant polycystic kidney disease (ADPKD) is the most common inherited renal disorder. Although a variety of candidate drugs have been found to modulate cystogenesis in animal studies, results from clinical trials have often been unfavorable due to low renal bioavailability and drug-induced side effects. To mitigate this, nanoparticles can be designed to deliver drugs directly to the target organ to increase effective dose while limiting off-target side effects. Unfortunately, there are no kidney-targeted nanomedicines clinically available, and most of the existing FDA-approved nanoparticles require intravenous administration which is not suitable for ADPKD that require lifelong therapy. To address this, we developed an oral drug delivery system using chitosan nanoparticles (CS-NP) that were loaded with peptide amphiphile micelles carrying metformin (met), an ADPKD drug candidate (CS-KM-met). We previously showed that CS-NP can shield met in the gastrointestinal tract; thus, we hypothesized that CS-NP could also enhance bioavailability of kidney-targeting micelles (KMs) upon oral administration. Specifically, we measured the loading capacity of KM-met in CS-NP, evaluated the stability of CS-KM-met under acidic conditions that mimic the gastric environment, and measured *in vitro* therapeutic effects. Upon oral administration in C57BL/6J mice, CS-KM-met showed significantly greater bioavailability and accumulation in the kidneys as compared to KM-met without CS-NP or free met for up to 24 hours. As such, CS-KM-met showed enhanced therapeutic efficacy *in vivo* upon oral administration in PKD mice (*Pkd1^fl/fl^; Pax8-rtTA; Tet-O-Cre*) compared to KM-met only. Herein, we demonstrate the potential of an oral delivery nanoformulation for the treatment of chronic kidney diseases such as ADPKD for the first time.

## 1. Introduction

Autosomal dominant polycystic kidney disease (ADPKD) is the most common monogenic chronic kidney disease (CKD), which affects 12.5 million people worldwide [1]. ADPKD is caused primarily by mutations in two genes, *PKD1* or *PKD2*, and is characterized by overproliferation of renal cells and uncontrolled cyst growth that leads to a loss in kidney function and eventual renal failure [2]. Tolvaptan, the only FDA-approved drug for ADPKD, is a vasopressin V2 receptor antagonist that inhibits adenylyl cyclase (AC) and the cyclic AMP (cAMP) pathway, decreases fluid secretion and cell proliferation, and thereby slows the progression of ADPKD. However, ADPKD patients undergoing tolvaptan treatment can experience significant side effects such as polyuria, thirst, nausea, and drug-induced liver damage [3–8].

In addition to tolvaptan, other drug candidates have shown promising therapeutic efficacy in preclinical studies, but enthusiasm was dampened during clinical trials due to severe side effects. For example, although bardoxolone methyl, a Nrf2 pathway activator that restores mitochondrial function and inhibits pro-inflammatory signals, showed benefits in reducing cystogenesis in ADPKD cell lines [9], bardoxolone methyl induced cardiac and gastrointestinal disorders in clinical trials [10]. Metformin (met), an AMPK activator and mTOR inhibitor, currently FDA approved for diabetes, was found to slow disease progression in recent phase 2 clinical trials in ADPKD patients [11, 12]. However, although mostly tolerable, off-target side effects were still found and included as hypoglycemia and lactic acidosis were observed [13–15]. Therefore, the development of drug delivery strategies such as kidney-targeting nanoparticles may benefit diseases like ADPKD by delivering drugs directly to the kidneys and simultaneously reducing off-target side effects.

To that end, our group previously reported on the development of peptide amphiphile micelles that were decorated with kidney-targeting peptides including (Lys–Lys–Glu–Glu–Glu)_3_–Lys ((KKEEE)_3_K), which binds megalin, a multiligand receptor highly expressed on renal tubular cells [16–21]. Although kidney-targeting micelles (KMs) effectively homed to the kidneys *in vivo* upon intravenous (IV) injection, IV administration is not practical for chronic diseases such as ADPKD that progress over a lifetime [22, 23].

To facilitate passage through oral administration and enhance bioavailability of drugs for ADPKD, our lab also reported on the development of chitosan nanoparticles (CS-NP) for loading ADPKD drugs and demonstrated improved therapeutic outcomes when formulated within CS-NP [24]. We chose to develop CS-NP because chitosan is biocompatible and a natural polysaccharide derived from crustacean shells. Importantly, chitosan has been used to improve the bioavailability of drugs using oral delivery due to its high stability in acidic conditions like that of the stomach and mucoadhesive properties that enable high absorption in the intestines [25–27].

Combining these previous works, in this study, we hypothesized that encapsulating KMs loaded with metformin into CS-NP would enable enhanced KM-met bioavailability and delivery to the kidneys upon oral delivery for ADPKD therapy. To test our hypothesis, we first incorporated metformin into KMs and tested the metformin release profile as well as the therapeutic effects of KM-met *in vitro.* Next, we evaluated the loading capacity and release of KM-met from CS-NP and studied the ability of CS-KM-met to cross an *in vitro* intestinal model and induce a therapeutic response on murine cortical collecting ducts cells (mpkCCD_c14_), or cell types most affected in ADPKD. Finally, we evaluated the pharmacokinetic properties of CS-KM-met *in vivo* and the therapeutic outcomes in an ADPKD mouse model. Overall, we present for the first time an oral delivery nanomedicine strategy designed for PKD and provides evidence for its practical implementation in chronic kidney diseases.

## 2. Methods and Materials

### 2.1 Synthesis of Therapeutic KMs

The (KKEEE)_3_K targeting peptide was synthesized using standard Fmoc-mediated solid phase peptide synthesis methods on rink Amide resin (Anaspec, Fremont, CA, USA) using an automated benchtop peptide synthesizer (PS3, Protein Technologies, Tucson, AZ, USA) as previously described [21]. A cysteine was added to the peptide sequence at the N-terminus to allow for a thioester linkage. Peptides were cleaved from the resin and deprotected with 94:2.5:2.5:1 by volume trifluoroacetic acid:1,2-ethanedithiol:H_2_O:triisopropylsilane and were precipitated and washed several times with cold diethyl ether, dissolved in water, lyophilized, and stored as powders at −20°C. Crude, peptide mixtures were purified by reverse-phase high performance liquid chromatography (HPLC) (Shimadzu, Kyoto, Japan) on a C8 column (Phenomenex, Torrance, CA, USA) at 50°C using 0.1% formic acid in acetonitrile/water mixtures and characterized by matrix-assisted laser desorption ionization time-of-flight (MALDI-TOF/TOF) mass spectral analysis (Autoflex Speed, Bruker, Billerica, MA, USA, Figure S1). Cysteine-containing peptides were conjugated to 1,2-distearoyl-sn-glycero-3-phosphoethanolamine-N-methoxy-poly(ethylene glycol 2000) (DSPE-PEG(2000)-maleimide, Avanti Polar Lipids, Alabaster, AL, USA) by adding an equimolar amount of the lipid to peptide in MilliQ water (pH 7.2). After gentle mixing for one week, the resulting DSPE-PEG(2000)-(KKEEE)_3_K was purified by HPLC on a C4 column as described above.

Fluorophore-conjugated amphiphiles were synthesized by conjugating Cy7 via a peptide bond to DSPE-PEG(2000)-amine (Avanti Polar Lipids, Alabaster, AL, USA) by adding an equimolar amount of Cyanine7 NHS ester (Lumiprobe, Hunt Valley, md, USA) to the lipid dissolved in 0.1 M aqueous sodium bicarbonate buffer (pH 8.5). After reaction at room temperature for over 24 hours, protected from ambient light, the mixture was purified on a C4 column and characterized as described above. Cy7-metformin was also synthesized via conjugating the ester group of Cy7 to the amine group of metformin hydrochloride as described above.

DSPE-PEG(2000)-metformin was synthesized by conjugating metformin hydrochloride via a peptide bond to DSPE-PEG(2000)-NHS (Avanti Polar Lipids, Alabaster, AL, USA) by adding a 5x molar excess of metformin to the lipid dissolved in 10 mM aqueous sodium carbonate buffer (pH 8.5). After reaction at room temperature for 24 hours, protected from ambient light, the mixture was purified on a C4 column and characterized as described above (Figure S1).

To self-assemble micelles, the appropriate DSPE-PEG(2000) amphiphiles were dissolved in methanol or chloroform and evaporated under a steady stream of air. The resulting film was dried under vacuum overnight and hydrated at 80°C with either MilliQ water or PBS, vortexed and sonicated as needed to obtain a clear solution and allowed to cool to room temperature. Therapeutic KM (KM-met Cy7) were composed of a monomer molar ratio of 10:45:45 consisting of DSPE-PEG(2000)-Cy7:DSPE-PEG(2000)-(KKEEE)_3_K:DSPE-PEG(2000)-metformin. NT-met Cy7 were composed of a monomer molar ratio of 10:45:45 consisting of DSPE-PEG(2000)-Cy7:DSPE-PEG(2000)-methoxy:DSPE-PEG(2000)-metformin.

### 2.2 Cell Culture

Mouse kidney cortical collecting duct (mpkCCD_c14_) cells were expanded in culture media comprised of DMEM/F12 supplemented with insulin, dexamethasone, selenium, transferrin, triiodothyronine, glutamine, d-glucose, epidermal growth factor (EGF), HEPES, sodium pyruvate as described [28]. Media was changed every two days and subcultures were passaged every 7-8 days. Human colon epithelial cells (Caco-2, ATCC HTB-37, ATCC, Manassas, VA, USA) were cultured following the manufacturer’s recommendations. Cells were expanded in Dulbecco’s Modified Eagle Medium (DMEM) supplemented with 10% fetal bovine serum (FBS) and 1% penicillin-streptomycin. *Pkd1* null proximal renal tubule cells (*Pkd1* null cells) isolated from *Pkd1flox/-:TSLargeT* mice were cultured in DMEM/F12 media, 2% FBS, 1x Insulin-Transferrin-Selenium (ITS-G), and ∼2 nM of tri-ido-sodium salt. Cells were expanded at 37 °C in a humidified incubator under 5% CO_2_. Cells at passage 3 were used for studies, and the media were changed every 2–3 days.

### 2.3 Drug Release of Metformin from KM-met

To assess the rate of metformin release from KM-met, 1000 µM of metformin-KMs in PBS was treated with proteases (3.3 U/mL from *streptomyces griseus*, Sigma Aldrich, St. Louis, MO, USA) and placed in Slide-A-Lyzer dialysis cassettes (Thermo Fisher Scientific, Waltham, MA, USA) with a molecular weight cutoff of 2000 Daltons in 10 mM sodium acetate buffer at pH 7.45. Metformin has a molecular weight of 129.16 g/mol, while DSPE-PEG(2000) and DSPE-PEG(2000)-metformin has a molecular weight of 2805 g/mol and 2895 g/mol, respectively, and are unable to escape the dialysis compartment. Measurements of free metformin release out of the dialysis chamber were made at select timepoints (30 min, 1, 2, 3, 4, 5, 6, 12 hrs) by UV-VIS spectrophotometer at an absorbance of 233 nm (Nanodrop, Thermo Fisher Scientific, Waltham, MA, USA).

### 2.4 *In vitro* cystogenesis with *Pkd1* null cells

50 μL of Matrigel^TM^ from BD Biosciences was added to each well in a 96 well plate and solidified in a 37° incubator for 15 minutes. *Pkd1* null cells were trypsinized and resuspended with 150uL of 2% Matrigel™ in assay medium to achieve approximately 3000 cells/well, and the cells were grown for 1–2 days. Cells were then treated with 300 µM metformin in KM-met, NT-met, or free met formulations, along with NT blank, (KKEEE)_3_K only, or PBS for 8 days. On day 8, cysts were imaged (Leica DMi8, Leica, Wetzlar, Germany), and the size of cysts was determined by ImageJ (NIH).

### 2.5 *In vitro* Therapeutic Effect of KM-met

To validate the therapeutic effect of KM-met, we treated polarized mice mpkCCD_c14_ cells on 0.4 µm pore Transwell membranes (Costar, Corning, NY, USA) with 300 µM of metformin, or an equivalent molar ratio of the therapeutic amphiphile within KM-met for 4 hours. Cultured cells were then lysed, and protein was extracted for Western blotting using standard protocols. Briefly, AMPK lysis buffer containing 1 mM dithiothreitol (Goldbio Technology, St. Louis, MO, USA), 1mM phenylmethylsulfonyl fluoride (Goldbio Technology) and 1X Complete™ Protease Inhibitor Cocktail (Sigma Aldrich, St. Louis, MO, USA) was used to lyse cells on ice with 50 μL of buffer per Transwell membrane. Cells were scraped with a cell scraper and kept on ice for 15 min. Samples were spun for 15 min at 14,000 RPM. After addition of sample buffer, samples were then incubated at 65°C for 15 min and immediately run on NuPAGE Bis-Tris gels (Thermo Fisher Scientific, Waltham, MA, USA) as per the manufacturer’s instructions. Proteins were transferred onto nitrocellulose membranes for immunoblotting and then probed with antibodies for phospho-AMPKα (Thr172) and β-Actin (Cell Signaling Technology, Danvers, MA, USA). The membrane was probed with appropriate secondary antibodies and imaged on an Odyssey Imaging System (LICOR, Lincoln, NE, USA).

### 2.6 Synthesis of CS-NP

The assembled micelles were loaded into CS-NP via ionic gelation [24]. Chitosan (Heppe Medical Chitosan GmbH, Halle, Germany) with 200 mPas viscosity and 85% degree of deacetylation was dissolved at a 2 mg/mL concentration in a solution of 0.5% acetic acid in MilliQ water. Then, a solution of 1 mg/ml anionic crosslinker of poly-L-glutamic acid sodium salt (Sigma Aldrich, St. Louis, MO, USA) was prepared. This crosslinker solution was added as the solvent to a thin film of KM-met, hydrating them and forming micelles. The chitosan solution was added dropwise under constant stirring to the crosslinker/micelle solution at a volume ratio of 5:2. An opalescent suspension was formed spontaneously. CS-NP were separated by microcentrifuging at 14,000 RPM at 14°C for 30 minutes. The pellet was then washed with increasing grades of ethanol in water and used immediately for studies, or frozen and lyophilized and stored at 4°–8°C.

### 2.7 Characterization of Micelle Loading into CS-NP

#### Dynamic Light Scattering (DLS)

Chitosan solution synthesized with 0-2000 μM of KM-met were filtered through Puradisc 0.2-μm polyvinylidene fluoride (PVDF) membrane filters (GE Healthcare Life Sciences, Pittsburgh, PA, USA) and measured immediately. DLS measurements were determined at 163.5° and 532 nm using a Wyatt Technology Möbiuζ system (Santa Barbara, CA, USA, N ≥ 3). All measurements were carried out at 25°C in MilliQ water after equilibrating for 5 minutes.

#### Transmission Electron Microscopy (TEM)

Negatively stained samples for TEM were prepared by placing NP synthesized with 0-2000 μM of KM-met in MilliQ water on 400 mesh lacey carbon grids (Ted Pella, Redding, CA, USA) for 5 minutes. Excess liquid was wicked away with filter paper and the grid was washed with MilliQ water before placing 2 wt.% uranyl acetate solution for 2 minutes, then washing with MilliQ water. Dried samples were immediately imaged on a JEOL JEM-2100F TEM (JEOL, Ltd., Tokyo, Japan).

### 2.8 Micelle Release from CS-NP and Morphological Response to pH

Micelle release studies from CS-NP was performed in simulated gastric fluid (SGF) composed of 2.0 g/L sodium chloride and 2.9 g/L HCl (pH 1.3), or simulated intestinal fluid (SIF) composed of 0.62 g/L sodium hydroxide and 6.8 g/L potassium phosphate monobasic (pH 6.8) [29]. KM-met released from CS-NP was quantified at 233 nm using a NanoDrop One microvolume UV-Vis spectrophotometer for up to 6 hours at room temperature (Thermofisher Scientific, Waltham, MA, USA). At the endpoint, the degraded CS-NP were spun down at 14,000 x g for 10 minutes, and the supernatant was measured in DLS to verify the presence of intact micelles.

### 2.9 *In vitro* Therapeutic Effect of CS-KM-met

To assess therapeutic efficacy of CS-KM-met *in vitro*, the cellular levels of phospho-AMPK (Kit #7959) and total AMPK (Kit #7961) were measured via enzyme-linked immunosorbent assays (ELISA, Cell Signaling Technologies, Danvers, MA, USA) according to the manufacturer’s instructions. All standards and samples were measured on a Varioskan LUX microplate reader at a wavelength of 450 nm. The phospho-AMPK to total AMPK ratio was normalized to the PBS group and presented in percentage.

### 2.10 Transepithelial Resistance (TER) measurement after CS-NP treatment

To test whether CS-KM-met can open tight junctions within the intestinal epithelium, Caco-2 cell monolayers were seeded onto Transwell inserts (Corning, NY, USA; diameter 6.5 mm, growth area 0.33 cm^2^, pore size 0.4 µm). At an initial density of 3 x · 10^5^ cell/cm^2^ and maintained for 21 days, a confluent monolayer was formed. The TER measurement was performed using an EVOM2 Epithelial Voltohmmeter (World Precision Instruments, USA). TER was measured 3, 2, and 1 days before treatment to establish baseline measurements, then every 6 hours immediately after treatment with CS-KM-met, CS-NP, free met, KM-met, or PBS control until 24 hours, and then again on days 2 and 3 post-administration.

### 2.11 Intravital Imaging of KM Micelles

To study the passage of KM through the glomerular filtration barrier (GFB), intravital imaging of KMs was performed. Under continuous anesthesia (isoflurane 1-2% inhalant via nose-cone), mice were placed on the stage of the inverted microscope with the exposed kidney mounted in a coverslip-bottomed chamber bathed in normal saline as described previously [30, 31]. Body temperature was maintained with a homeothermic blanket system (Harvard Apparatus, Holliston, MA, USA). Alexa Fluor 488-conjugated Dextran 500kDa (Thermo Fisher, Waltham, MA, USA) was administered intravenously by retro-orbital injections to label the circulating plasma (30 µL intravenous (IV) bolus from 10 µg/ml stock solution). 100 µL of 30 µM Cy7 labeled KMs or 3 µM disassembled monomers ((KKEEE)_3_K-Cy7-DSPE-PEG(2000)) were administered into the canulated carotid artery. The volume of blood in a mouse is ∼1.5-2 mL and the concentration of monomers (3 µM) in circulation will then be approximately 0.15-0.2 µM, which is below the critical micelle concentration of ∼1 µM [32]. The images were acquired using a Leica SP8 DIVE multiphoton confocal fluorescence imaging system with a 40× Leica water-immersion objective (numerical aperture (NA) 1.2) powered by a Chameleon Discovery laser at 960 or 1100 nm (Coherent, Santa Clara, CA) and a DMI8 inverted microscope’s external Leica 4Tune spectral hybrid detectors (emission at 510-530 nm for GFP, at 580-640 nm for AF594, and at 720-850 for Cy7) (Leica Microsystems, Heidelberg, Germany). The potential toxicity of laser excitation and fluorescence to the cells was minimized by using a low laser power and high scan speeds to keep total laser exposure as minimal as possible. Fluorescence images were collected in volume and time series (xyt, 526 ms per frame) with the Leica LAS X imaging software and using the same instrument settings (laser power, offset, gain of both detector channels).

### 2.12 Biodistribution of Orally Administered Cy7-labeled CS-KM-met

To evaluate the biodistribution of CS-KM-met, 6-7-week-old male and female C57BL/6J mice (Jackson Laboratories, Bar Harbor, ME, USA) were orally gavaged with 200 μL of 500 µM Cy7-labeled CS-KM-met, CS-NT-met, KM-met, NT-met, or free met. Mice were euthanized 24 hours post-injection and organs (e.g., brain, heart, lungs, liver, kidneys, spleen, intestines, and bladder) were excised and imaged *ex vivo* on an AMI HTX *in vivo* imaging system (Spectral Instruments Imaging, Tuscon, AZ, USA). The fluorescence signal was quantified via Aura software (Spectral Instruments Imaging, Tucson, AZ, USA, n ≥ 4), and background was subtracted from the PBS-treated group. The mean radiance (photons/s/cm^2^/sr) for each organ was quantified as a region of interest, and % of total organ fluorescence was obtained by dividing each organ by the sum of all the organ regions. Blood draws were performed either retro-orbitally or via tail vein at 30 min, 3, 6, 12, and 24 hrs post-administration. Fluorescence was measured in serum and quantified using a Cy7 met calibration curve developed in mouse serum. All animal procedures followed NIH guidelines for the care and use of laboratory animals and were approved by the University of Southern California’s Institutional Animal Care and Use Committee.

### 2.13 Therapeutic Efficacy in ADPKD Mice

To assess the ability of CS-NP to enhance the therapeutic efficacy of orally administered KM-met in ADPKD, CS-KM-met or KM-met (300 mg/kg met) was administered in *Pkd1^fl/fl^; Pax8 rtTA;Tet-O-Cre* mice. To induce a slowly developing ADPKD model mimicking the human condition, mice were IP injected with doxycycline (50 mg/kg/day) on postnatal day 27-29 (P27-P29), and again on P43 and P57. Mice were orally gavaged every three days starting on P30 and euthanized on P120. Kidneys were excised to assess kidney weight to body weight (KW/BW) ratio and H&E stained to measure cystic index. Cystic index was defined as the percentage of cystic area divided by total kidney area [33] and measured by ImageJ.

### 2.14 Kidney Health in ADPKD Mice

Kidney health markers including measurements of plasma sodium (Na), potassium (K), chloride (Cl), ionized calcium (iCa), total carbon dioxide (tCO2), glucose (Glu), blood urea nitrogen (BUN)/Urea, creatinine (Crea), hematocrit (Hct), and hemoglobin (Hb) were assessed. On the day of harvest, 90 µL of blood taken from the submandibular vein was analyzed using Chem-8+ cartridges for the i-Stat Handheld Blood Analyzer (Abbott, Chicago, IL, USA).

### 2.15 Statistical Analysis

A Student’s *t*-test was used to compare means of pairs using GraphPad Prism 8 (San Diego, CA). Analysis of variance (ANOVA) with a Tukey’s test for post-hoc analysis was used to determine statistical significance, and *p* ≤ 0.05 was considered to be significant.

## 3. Results and discussion

### 3.1 Synthesis, Characterization, and *in vitro* Therapeutic Effects of KM-met

To first construct kidney-targeting micelles (KMs), the kidney-targeting peptide (KKEEE)_3_K, metformin, and the fluorophore Cy7 was conjugated to DSPE-PEG(2000) and self-assembled under aqueous conditions (Figure 1a) [34]. To characterize the size and shape of KM-met, we imaged KM-met by TEM and found the particles were monodispersed and of spherical morphology (Figure 1b). Next, to verify that free metformin can be released from micelles intracellularly, a commercially available cocktail of proteases derived from *streptomyces griseus* were incubated with KM-met [35]. After 12 hours of incubation, 35.7 ± 3.2 % metformin was released from KM, while no appreciable drug release was found in the absence of proteases (Figure 1c), confirming that the peptide bond between DSPE-PEG(2000) and metformin was cleaved by proteases as expected [36].

**Figure 1.**
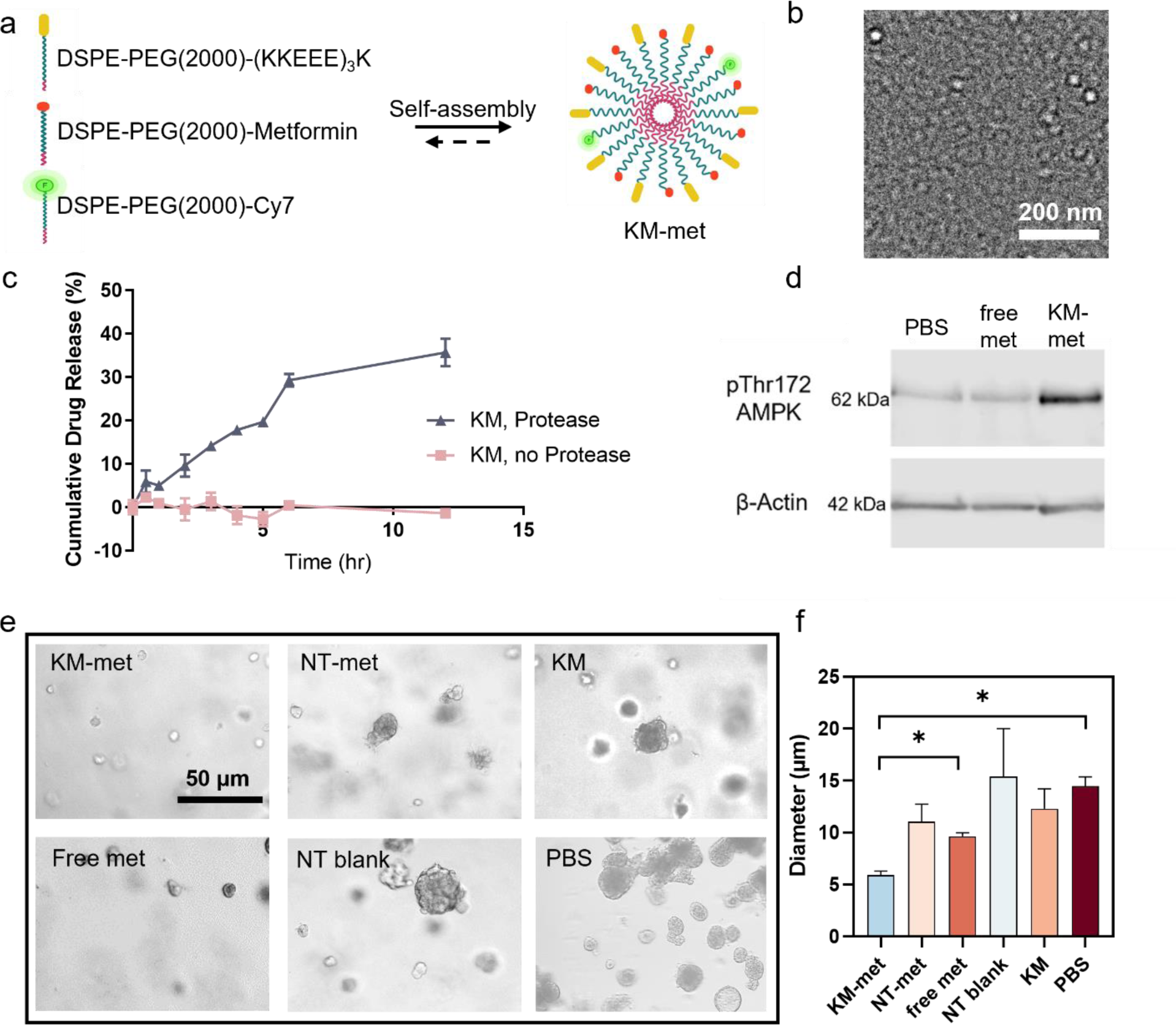
Characterization and *in vitro* therapeutic effects of KM-met. (a) Schematic of KM-met self-assembly. (b) TEM confirms spherical morphology and the size of KM-met to be approx. 15 nm. (c) Drug release of metformin from KM over time via protease cleavage. (d) Western blot shows higher levels of pThr172 AMPK in mpkCCD_c14_ cells when treated with KM-met compared to free metformin (at 300 µM) or PBS for 4 hours. (e, f) Brightfield images and quantification of cystogenesis using *Pkd1* null cells treated with KM-met, free met, NT-met, NT blank, KM, or PBS (*p ≤ 0.05, N ≥ 4).

Initial therapeutic effects of metformin released from KMs was evaluated *in vitro* through activation of the AMPK pathway in mpkCCD_c14_ cells by Western blotting. After 4 hours of treatment at an equivalent concentration of 300 µM, higher levels of the activated form of AMPK, pThr172 AMPK, were observed in cells treated with KM-met as compared to free metformin or PBS after 4 hours (relative intensity: 1.20 KM-met vs. 0.39 free met vs. 0.30 PBS, Figure 1d). In addition, the *in vitro* therapeutic effects of KM-met was assessed *via* a 3D cyst model (Figure 1e). Specifically, *Pkd1* null cells generated from the *Pkd1flox/-:TSLargeT* mouse model of ADPKD were cultured in Matrigel^TM^ and treated with KM-met, NT-met, or free met at a metformin concentration of 300 µM and the effects on cyst growth were measured on day 8. Empty non-targeting (NT) micelles, KM, and PBS without metformin also served as additional controls. Quantification of the cyst images showed that the KM-met treated group had significantly smaller cyst diameter (5.9 ± 0.9 µm) when compared to free met (9.6 ± 0.5 µm, *p* ≤ 0.05) and PBS treated group (9.6 ± 0.5 µm, *p* ≤ 0.05), which confirms that the targeting ability of KMs enables enhanced drug delivery of met and an increase in therapeutic effects *in vitro* (Figure 1f).

### 3.2 Loading of KMs within CS-NP

To load KM-met into CS-NP, we first mixed KM-met into the anionic poly-L-glutamic acid crosslinker solution. The chitosan solution was then added to the crosslinker/micelle solution to form CS-KM-met through ionic gelation [37–39]. KM-met was not mixed with the chitosan solution because the solution became too viscous and could not be used for CS-NPs formation.

CS-NPs loaded with KM-met were found to be ∼160 nm in diameter using dynamic light scattering (DLS). As found in Figure 2a and b, KMs on their own are 13.9 ± 1.8 nm in diameter and CS-NPs are 155.9 ± 14.8 nm as measured by DLS. When micelles were mixed with CS-NPs separately, two distinct size distributions (15.0 ± 2.1 nm and 148.2 ± 17.9 nm) can be observed, which correspond to the expected micelle size of ∼14 nm and the CS-NPs of ∼160 nm (Figure 2c). However, when micelles are loaded into CS-NPs, the micelle only peak of ∼14 nm is no longer observed, confirming that KMs are successfully loaded into CS-NPs (Figure 2d).

**Figure 2.**
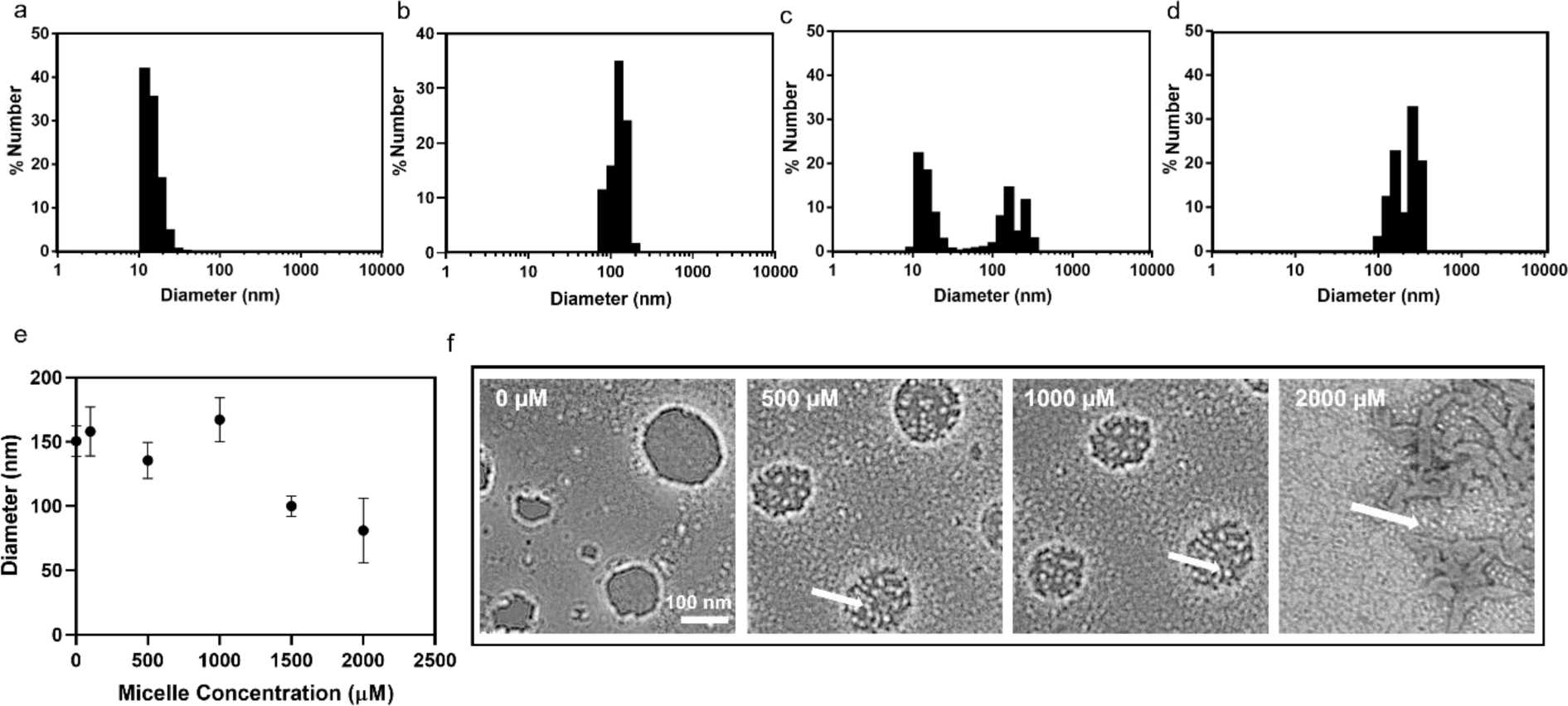
KM loading into CS-NP. Representative DLS measurements of (a) KM, (b) CS-NP, (c) KM mixed with CS-NP, and (d) KM loaded into CS-NP. Micelle peaks corresponding to ∼14 nm were present in the mixed condition (c) but were not seen in the loaded condition (d). (e) DLS measurements of CS-NP loaded with various initial starting concentrations of KM. (f) TEM images of CS-NP synthesized with varying concentrations of micelles (arrows) between 0 µM and 2000 µM.

Next, we probed the maximum micelle concentration that could be used to synthesize micelle-loaded CS-NP. As found in Figure 2e, micelle concentrations added to the crosslinker solution of up to 1000 µM had similar sizes to that of unloaded CS-NP (∼160 nm). Beyond 1000 µM micelle concentration, the diameter of the CS-NP decreases while polydispersity increases, showing a loss of stability. As also confirmed by TEM in Figure 2f, individual micelles are found encased within CS-NP up to 1000 µM KM-met. However, at 2000 µM KM-met, CS-NP are no longer present and, instead an unordered structure with the majority of micelles existing separately from the bulk chitosan material is observed (arrows, Figure 2f). Thus, we concluded that a maximum of 1000 µM micelles in the crosslinker solution can be combined with the chitosan solution for synthesis of micelle-loaded CS-NP.

### 3.3 KM-met Release from CS-NP and CS-KM-met Penetration and Therapeutic Effects Using *in vitro* Intestinal and Kidney Models

As previously reported, CS-NP are stable in low pH but can degrade under neutral pH similar to the intestinal environment to enable drug release and absorption into the bloodstream upon oral delivery [24]. To test the stability of CS-KM-met under low a pH environment as in the stomach, we incubated CS-KM-met in simulated gastric fluid (SGF, pH = 1.3). After 6 hours, 29.1 ± 2.5% of total KM-met was found to be released, as measured by the metformin wavelength of 233 nm. This is in contrast to CS-KM-met incubation in simulated intestinal fluid (SIF, pH = 6.8) which released 70.6 ± 1.5% of KM-met (p ≤ 0.01) (Figure 3a). To validate whether the released KM-met from CS-NP under neutral pH were intact micelles, or disassembled monomers, we measured the size of samples at the final timepoint and found intact micelles of ∼14 nm size were released into the supernatant (Figure 3b).

**Figure 3.**
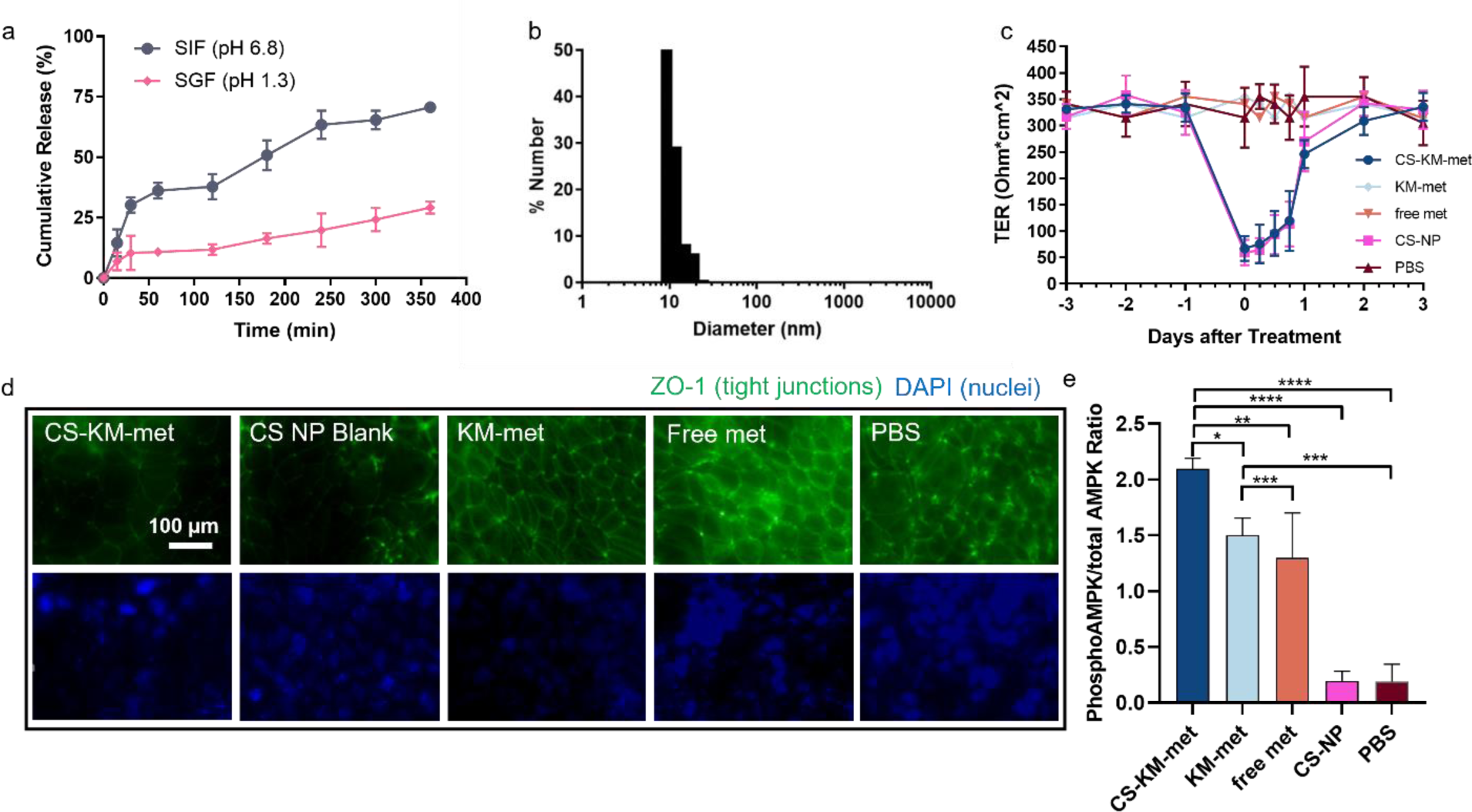
*In vitro* therapeutic efficacy of CS-KM-met. (a) *In vitro* release of KM from CS-KM under pH conditions found in the GI tract (pH = 1.3, SGF; 6.8 SIF, N ≥ 4). (b) DLS size measurements of micelles released from SIF at 3 hr. (c) TER measurements of Caco-2 monolayers incubated with CS-KM-met, KM-met, free met, CS-NP, or PBS for up to 6 days. (d) Caco-2 monolayers stained with ZO-1 and DAPI at 6 hr. (e) Phosphorylated AMPK to total AMPK normalized to PBS group obtained *via* ELISA on mpkCCD_c14_ cells to test the therapeutic efficacy of CS-KM-met (*p ≤ 0.05, **p ≤ 0.01, ***p ≤ 0.001, ****p ≤ 0.0001, N ≥ 3).

In addition to protection against an acidic environment mimicking the stomach, chitosan has been reported to increase bioavailability by opening tight junctions within the intestinal epithelium [40]. To verify this effect, transepithelial resistance (TER) was measured on human colorectal Caco-2 cells cultured on Transwell permeable supports following treatment with CS-KM-met [41, 42]. TER was measured three days before treatment, and again after administration of nanoparticles or free drug for up to 3 days. Within 6 hours post-administration of CS-KM-met, a significant 79.9% reduction in resistance was observed (from 334.0 ± 26.8 Ohm*cm^2^ to 67.1 ± 23.3 Ohm*cm^2^). On the other hand, no changes in TER were observed for samples treated with KM-met, free met, or PBS (Figure 3c) [24]. Notably, a recovery to pre-treated baseline resistance levels (335 Ohm*cm^2^) was observed at 3 days post-treatment, demonstrating that these effects on tight junctions are transient [43]. Additionally, we observed a decrease in ZO-1 signal in treatment groups that contained chitosan at 6 hours after treatment (Figure 3d), suggesting that CS-KM-met can enhance penetration through the intestinal epithelium similar to our previous reports [24]. Overall, these results using *in vitro* models show that CS-KM-met has the potential to protect cargo under acidic conditions like the environment of the stomach, enhance paracellular transport across the intestinal epithelium, and enable micelle-drug release at a neutral pH similar to the intestinal environment, thereby promoting higher absorption and bioavailability in circulation [44].

To test the delivery and therapeutic efficacy of CS-NP-met, we utilized a simple Transwell model that was seeded with Caco-2 intestinal cells on the apical side of the Transwell membrane and mpkCCD_c14_ cells grown on the bottom of the plate. After 12 hours, we found that mpkCCD_c14_ cells in the CS-KM-met treatment group showed the highest phosphorylated AMPK to total AMPK ratio after normalizing to PBS group, as found through ELISA assays, among all met-containing groups: 6.9 ± 2.1 % for free met, 7.9 ± 0.8 % for KM-met, and 11.1 ± 0.5 % for CS-KM-met (*p* ≤ 0.005), while no change was found upon CS-NP treatment (normalized to PBS). This confirms that the *in vitro* delivery of metformin and activation of AMPK can be improved through the combined chitosan and kidney targeting platforms and is the greatest in CS-KM-met group as compared to all other groups (Figure 3e).

### 3.4 *In vivo* Biodistribution of CS-KM-met

To evaluate the ability of CS-NP to enhance micelle bioavailability *via* oral delivery *in vivo*, wildtype C57BL/6J mice were orally administered Cy7-labeled CS-KM-met. Over 24 hours, CS-KM-met treated mice consistently showed higher serum fluorescence as compared to CS-NT-met, KM-met, NT-met and free met, demonstrating enhanced bioavailability and depot into systemic circulation (Figure 4a). Upon *ex vivo* imaging and analysis, Cy7-labeled CS-KM-met also showed higher kidney accumulation (28.9 ± 4.1 %) when compared to CS-NT-met (20.9 ± 4.1 %, *p* ≤ 0.01) (Figure 4b). Similarly, Cy7-labeled KM-met demonstrated higher renal accumulation (15.9 ± 2.1 %) when compared to NT-met (8.7 ± 0.6 %, *p* ≤ 0.01), confirming the benefits of CS-NP in enhancing bioavailability through oral administration and the renal targeting ability through the (KKEEE)_3_K peptide *in vivo*.

**Figure 4.**
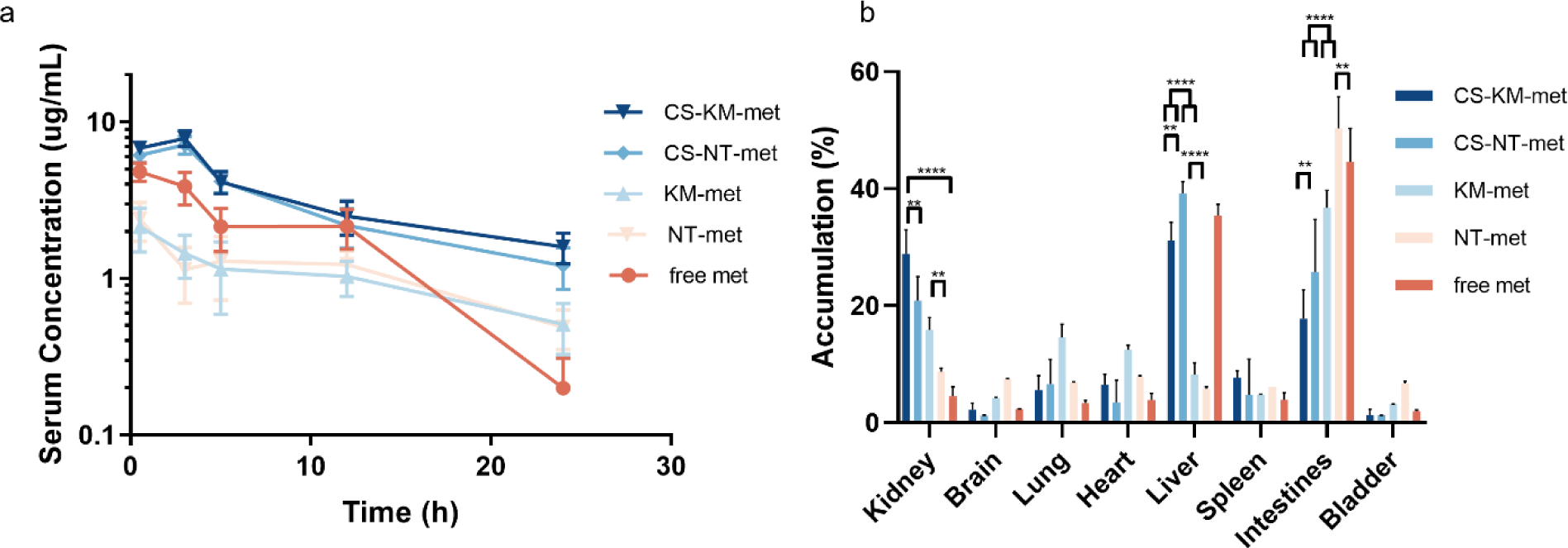
Bioavailability and biodistribution of Cy7-labeled CS-KM-met 24 hours after oral gavage. (a) Serum levels of Cy7-labeled micelles in CS-KM-met is the highest after oral delivery. (b) Quantification of *ex vivo* organ Cy7 fluorescence levels showed higher accumulation in the kidneys for Cy7-labeled CS-KM-met and CS-NT-met (**p ≤ 0.01, ****p ≤ 0.0001, N = 4).

In addition, intravital imaging, which provides real-time imaging of glomerular filtration and renal cell uptake, was performed to evaluate the mechanism by which KMs enter the kidneys. Consistent with our previous studies [18], Figure S3 and Video S1 shows Cy7-micelles enter the glomerulus, pass the glomerular filtration barrier (GFB), and access the Bowman’s space and tubules upon administration via the carotid artery. Interestingly, free amphiphilic monomers only entered the glomerulus but did not pass the GFB or enter the Bowman’s space (Figure S2, Video S2). This may be due to monomers binding to serum components such as albumin, which do not normally pass enter the GFB [45, 46]. In contrast, zwitterionic properties of the megalin-binding peptide, (KKEEE)_3_K, in addition to PEGylation, hinders serum protein adsorption of KMs [18]. Thus, in general, these results show that after oral delivery and depot into the bloodstream, KMs are able to passage through the GFB and access the renal tubular system, which is highly applicable for drug delivery in ADPKD.

### 3.5 Therapeutic Efficacy of CS-KM-met in ADPKD Mice

The therapeutic effects of CS-KM-met upon oral administration was tested in a slowly progressing ADPKD murine model using *Pkd1^fl/fl^;Pax8-rtTA;Tet-O-Cre* mice [47]. In this model, a slowly progressing PKD phenotype is developed by conditional knockout of the *Pkd1* gene, induced by doxycycline injection. After induction of *Pkd1* knockout starting with IP doxycycline injections at postnatal day 27 (P27), mice were orally administered every three days CS-KM-met or KM-met with an oral metformin dose of 300 mg/kg/day (Figure 5a), as reported previously in ADPKD preclinical studies [13, 24, 48], and euthanized on P120 when severe disease is expected using this model [49]. As found in Figure 5, mice treated with CS-KM-met showed a decreased cystic index (i.e., the percentage of cystic area divided by total kidney area) as compared to KM-met, (9.6 ± 4.2% vs. 24.6 ± 12.0 %, *ns*), as well as the no treatment control group (9.6 ± 4.2% vs. 41.5 ± 8.8%, *p* ≤ 0.05, Figure 5b, c). Importantly, CS-KM-met-treated mice also showed a lower kidney weight/body weight (KW/BW) ratio (1.3 ± 0.3%) as compared to KM-met group (2.3 ± 2.0%, *ns*) and the no treatment control group (5.4 ± 0.6 %, *p* ≤ 0.05), (Figure 5d). Nonetheless, kidney health markers including blood urea nitrogen (BUN) and creatinine were found to remain similar between CS-KM-met, KM-met and control groups, indicating that CS-KM-met has high biocompatibility and minimal toxicity (Table 1). In summary, CS-KM-met showed higher therapeutic efficacy in ADPKD mice as compared to KM-met and also demonstrated high biocompatibility profile. To our knowledge, KMs loaded into CS-NP is the first development of an oral delivery nanoformulation for kidney disease.

**Figure 5.**
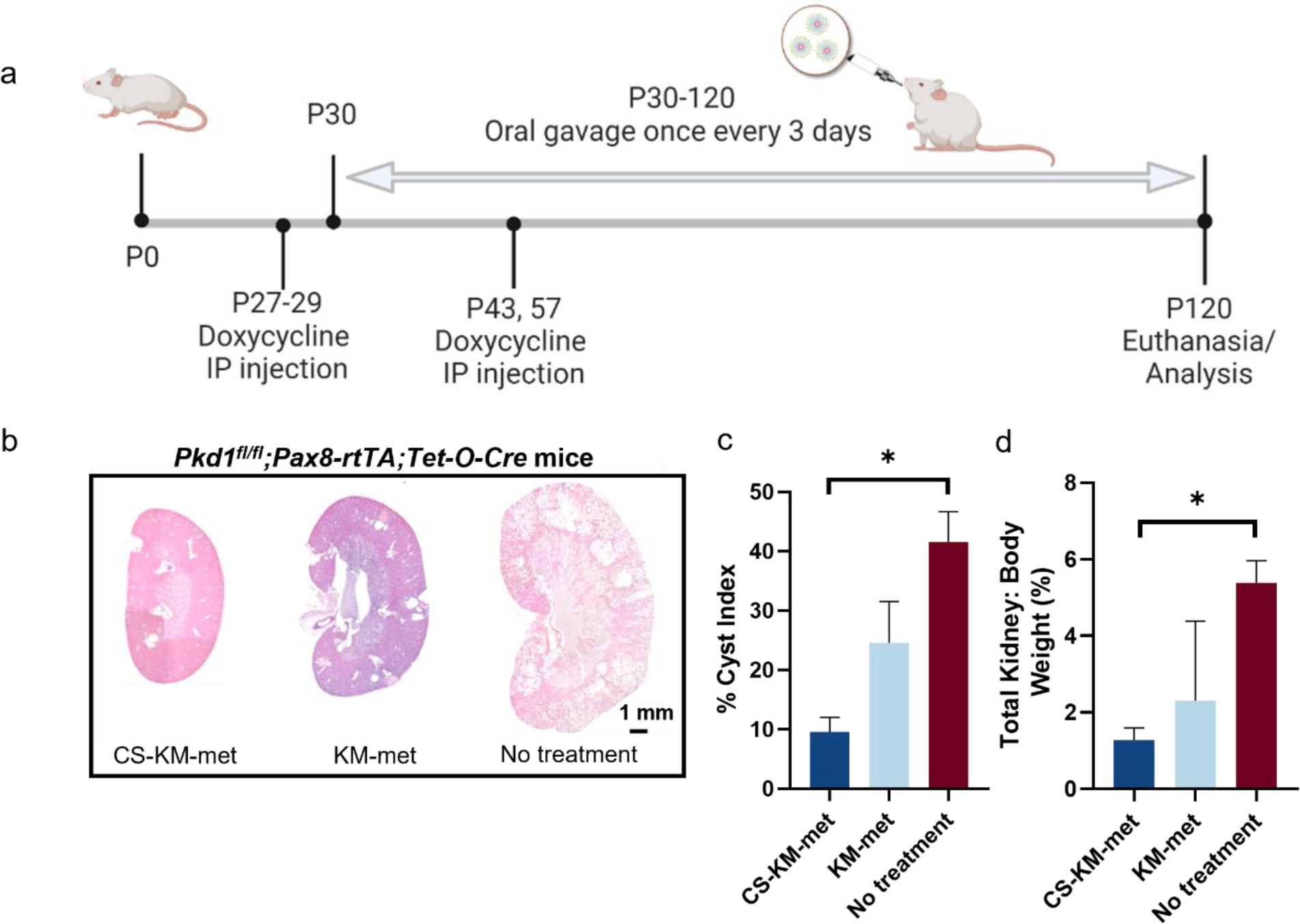
Therapeutic efficacy of orally administered CS-KM-met in ADPKD mice. (a) H&E staining of whole kidneys shows smaller kidneys with fewer cysts in the CS-KM-met treatment group. (b) Cystic index of the CS-KM-met treated group is significantly decreased as compared to KM-met and no treatment groups. (c) A lower KW/BW ratio was also found in the CS-KM-met treated group (*p ≤ 0.05, N = 3).

**Table 1.**
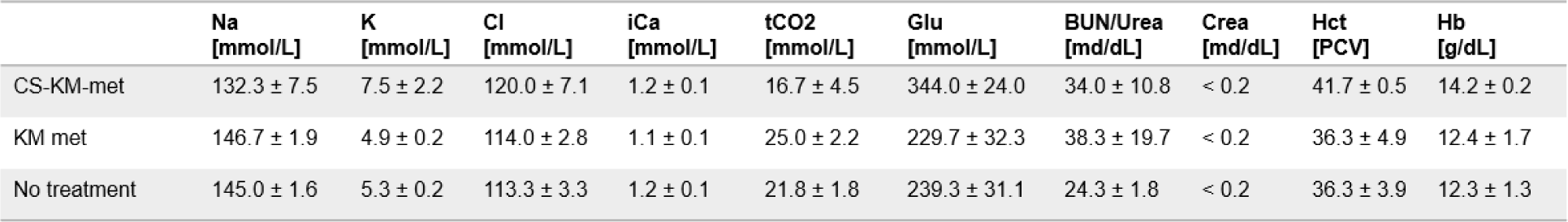
Serum components, electrolytes, and kidney health markers of ADPKD mice.

## 4. Conclusion

In this study, we report a novel, combinatorial oral drug delivery strategy for ADPKD using CS-NP containing KM-met (CS-KM-met). By incorporating KM-met into CS-NP, CS-KM-met were able to overcome the physiological barriers such as low pH in the stomach and enable high bioavailability of KM-met in systemic circulation. As such, CS-KM-met-treated mice demonstrated the highest renal accumulation of Cy7-label micelles for up to 24 hours. Upon met delivery using CS-KM-met via oral gavage in a murine model of ADPKD, cystic index and total kidney/body weight ratio of mice were significantly reduced compared to KM-met and control groups, demonstrating CS-KM-met provides a safe and long-term method for delivering drugs by oral administration with repeated doses for chronic diseases such as ADPKD. Although our study demonstrates improved therapeutic outcomes of metformin, future research will focus on loading other drugs that have higher toxicity profiles such as tolvaptan, which can cause drug-induced liver toxicity. In addition, future studies will test the efficacy of our KM-drug/CS-NP system in other PKD models, given that there are over 400 pathogenic *PKD1* mutations in patients with ADPKD [50] and in addition to *PKD1* mutations, ADPKD also includes *PKD2* mutations in 15% of patients. Nonetheless, our studies show the therapeutic potential and safety of KM-drugs loaded into CS-NP as a strategy for oral drug delivery in ADPKD, and the insights gained from this work can be more broadly applied to other chronic kidney diseases.

## Acknowledgements

The authors would like to acknowledge the financial support from the University of Southern California (USC) Alfred E. Mann Institute (AMI) fellowship awarded to J.W., and the New Innovator Award (NIH, DP2-DK121328), NSF EAGER from DMR BMAT 2132744, and WISE Major Support Award granted to E.J.C. The authors would also like to thank the Center for Electron Microscopy and Microanalysis (CNI) at USC for assistance in TEM imaging. In addition. fluorescence imaging was performed at the USC Multi-Photon Microscopy Core funded by the National Institute of Health grant S10OD021833.

## Supplementary Information

**Figure S1.**
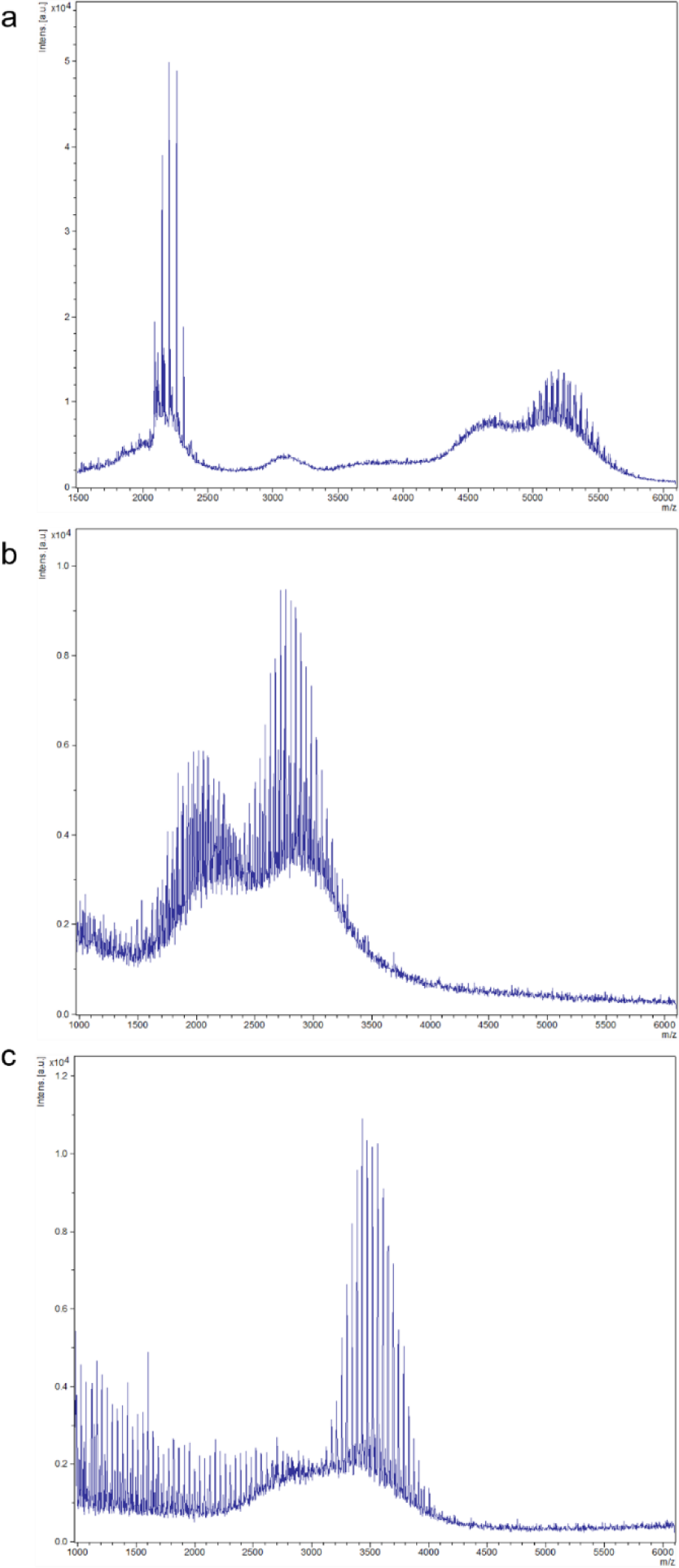
MALDI characterization of (a) DSPE-PEG(2000)-(KKEEE)_3_K (expected m/z: 5171 g/mol), (b) DSPE-PEG(2000)-Metformin (expected m/z 2895 g/mol), and (c) DSPE-PEG(2000)-Cy7 (expected m/z: 3522 g/mol).

**Figure S2.**
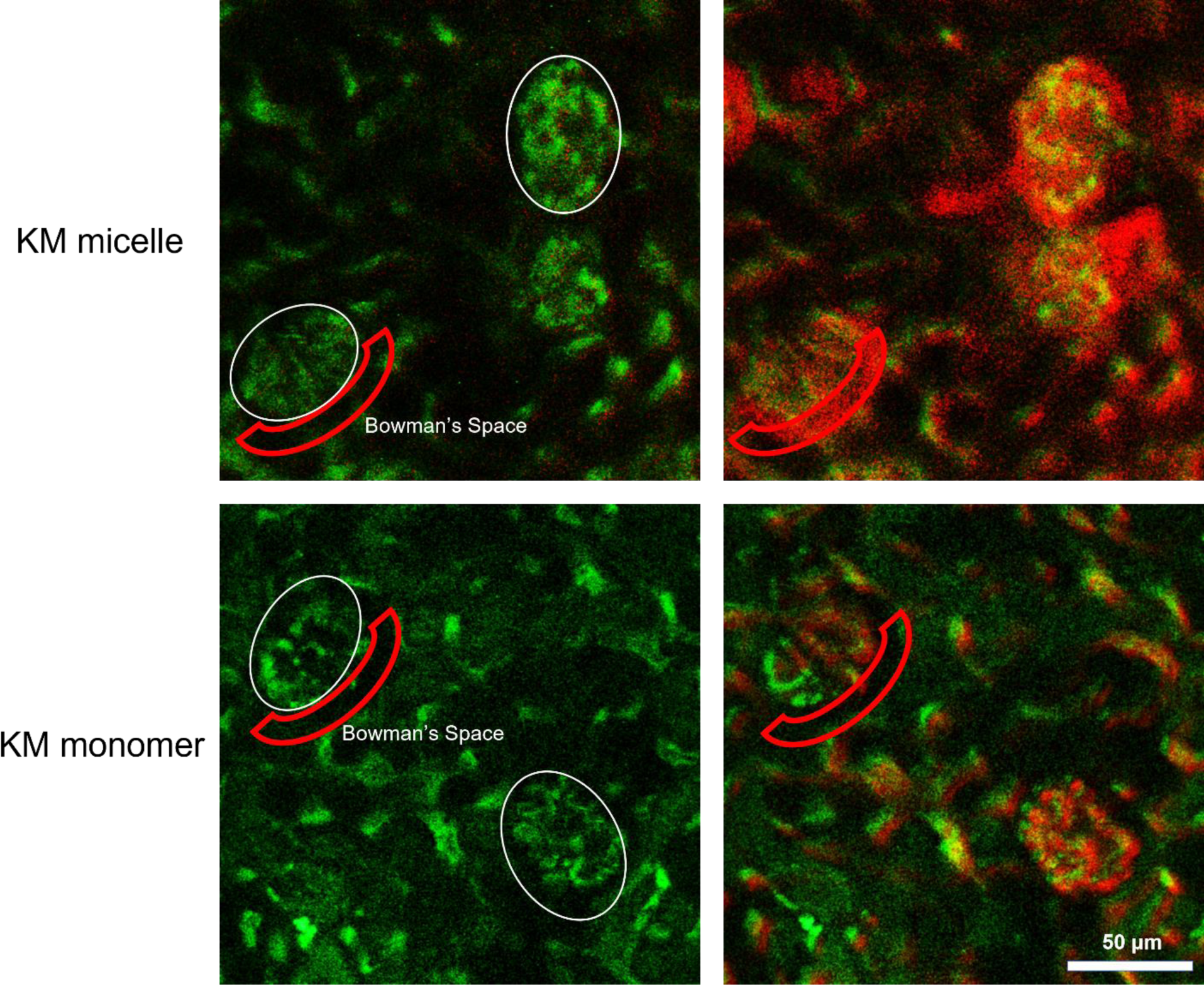
Intravital images of kidney glomeruli (white circle) after KM micelles and KM monomers were injected into the carotid artery of C57BL/6 mice. Both KM micelles and KM monomers entered the glomerulus, but only KM micelles were able to pass through the GFB and enter the Bowman’s space. 500kDa Dextran-Alexa Fluor 488 (green, plasma), Cy7-labeled KMs (red).

Video S1. Intravital images of KM micelles injected into the canulated carotid artery of C57BL/6 mice.

Video S2. Intravital images of KM monomers injected into the canulated carotid artery of C57BL/6 mice.

