## Supplemental file for "Oral Delivery of Kidney Targeting Nanotherapeutics for Polycystic Kidney Disease"

*Corresponding author: Eun Ji Chung

Supplementary Information

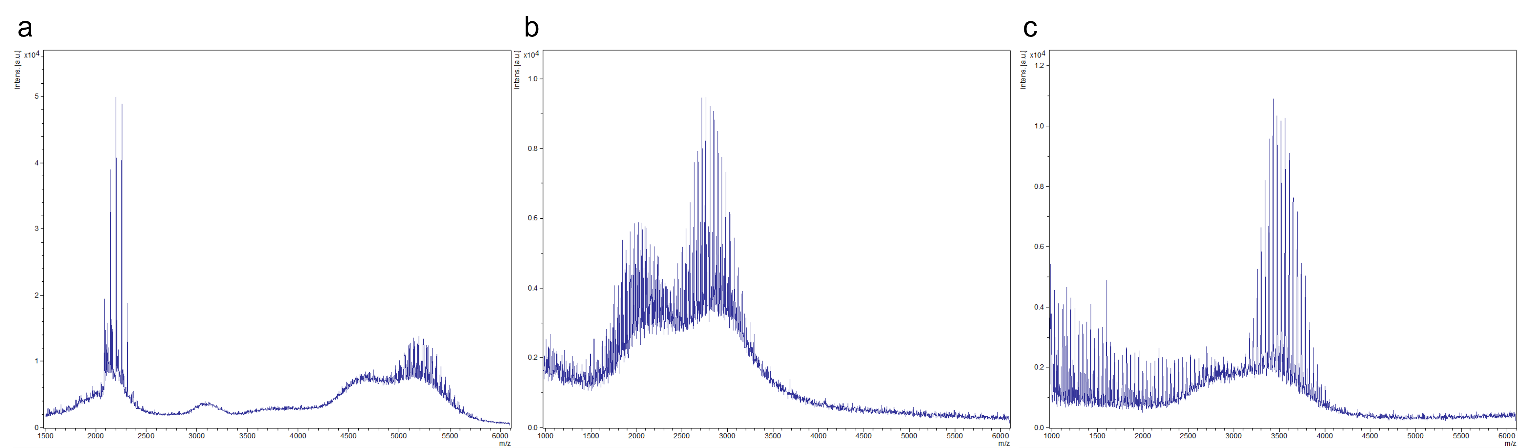

Figure S1. MALDI characterization of (a) DSPE-PEG(2000)-(KKEEE)_3_K (expected m/z: 5171 g/mol), (b) DSPE-PEG(2000)-Metformin (expected m/z 2895 g/mol), and (c) DSPE-PEG(2000)-Cy7 (expected m/z: 3522 g/mol).

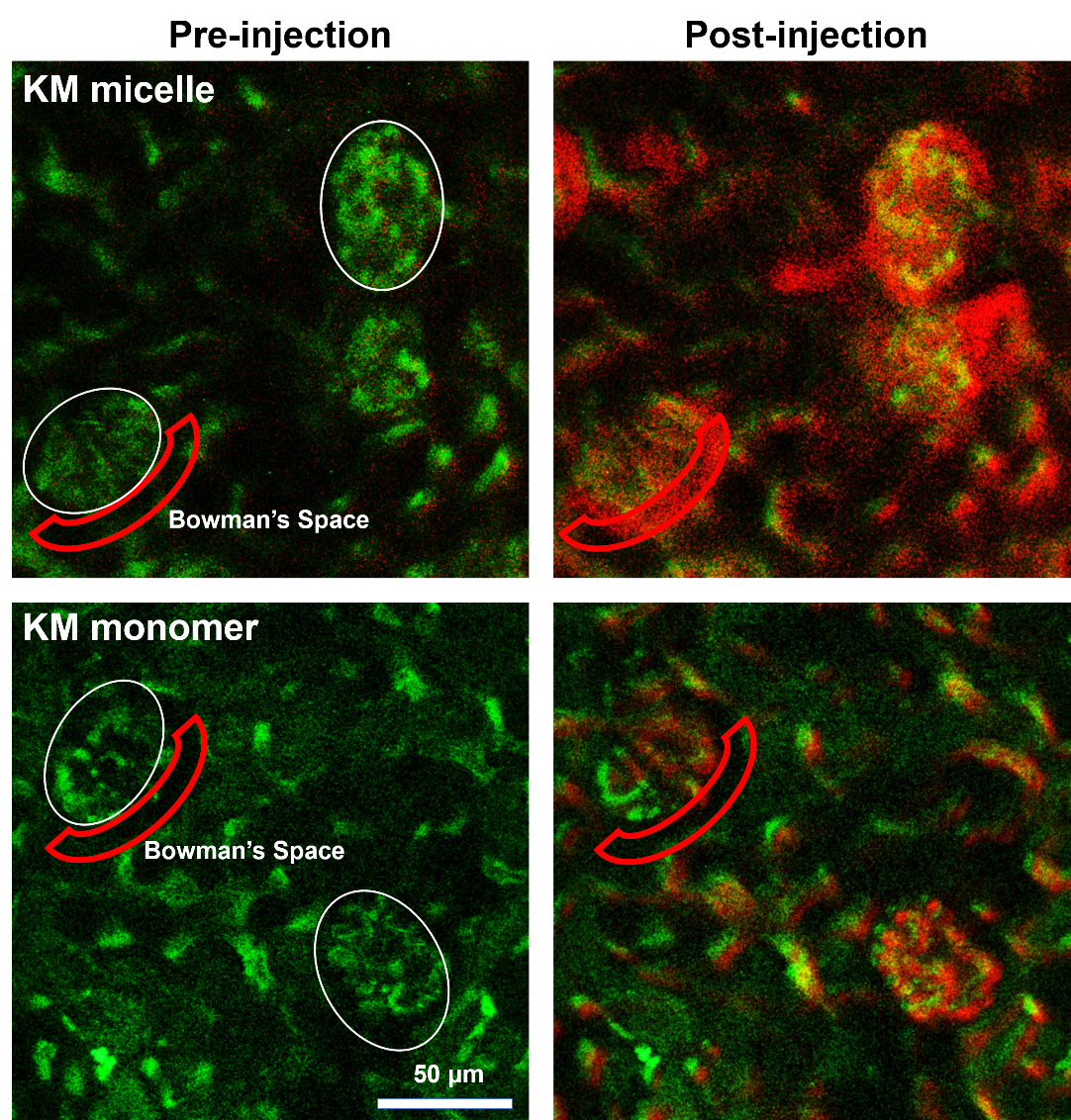

Figure S2. Intravital images of kidney glomeruli (white circle) after KM micelles and KM monomers were injected into the carotid artery of C57BL/6 mice. Both KM micelles and KM monomers entered the glomerulus, but only KM micelles were able to pass through the GFB and enter the Bowman’s space. 500kDa Dextran-Alexa Fluor 488 (green, plasma), Cy7-labeled KMs (red).

Video S1. Intravital images of KM micelles injected into the canulated carotid artery of C57BL/6 mice.

Video S2. Intravital images of KM monomers injected into the canulated carotid artery of C57BL/6 mice.
